# Exploring the Interaction of APOE-*ε*4 and PICALM rs3851179 With Dynamic Functional Connectivity in Healthy Middle-Aged Adults at Risk for Alzheimer’s Disease

**DOI:** 10.64898/2025.12.16.693545

**Authors:** Shyamal Y. Dharia, Camilo E. Valderrama, Qian Liu, Stephen D. Smith

## Abstract

**Objective:** This study investigates whether dynamic functional connectivity (dFC) dwell-time patterns derived from resting-state fMRI (rs-fMRI) can distinguish Alzheimer’s disease (AD) genetic risk profiles, specifically the *APOE*-*ε*4 (A+) and *PICALM* rs3851179 (P+) variants, in cognitively healthy, middle-aged adults.

**Approach:** We estimated recurring dFC clusters from rs-fMRI data and quantified the dwell-time (total duration spent in specific connectivity states) for three cohorts: not-at-risk, A+P-, and A+P+. To evaluate the utility of these temporal features, group differences in dwell-time profiles were assessed, and logistic regression with permutation testing was employed to classify genotypes based on dFC patterns.

**Main results:** Individuals in at-risk groups (A+P- and A+P+) exhibited significantly reduced dwell-time in left-hemisphere hubs compared to the not-at-risk group, aligning with known left-hemisphere vulnerability in early AD progression. The logistic regression models achieved above-chance discrimination of genotypes, with permutation tests confirming a significant trend when distinguishing not-at-risk individuals from the combined at-risk cohorts.

**Significance:** These findings suggest that the temporal dFC features are sensitive to subtle functional brain alterations linked to AD genetic risk before clinical symptoms appear. Dwell-time features represent a promising physiological marker for early risk stratification and warrant further validation in larger longitudinal datasets. Our code is available at https://github.com/Shyamal-Dharia/APOE-PICALM-dFC-dwell-time.git.

## Introduction

Alzheimer’s disease (AD) is a neurodegenerative disorder involving progressive memory loss and the deterioration of executive functions (Scharre, 2019; Rao et al., 2022). Although symptoms of AD are typically reported after the age of 65, psychometric tests can sometimes detect subtle declines in cognitive abilities and social engagement up to a decade earlier (Langbaum et al., 2020). Such results suggest that there are potential behavior patterns and biomarkers that can predict the likelihood that an individual will develop AD (see Breijyeh and Karaman (2020), for a review). The challenge for researchers is to identify these markers in order to provide patients and clinicians with the earliest possible indicator of potential AD so that physicians can prescribe medications that could slow the progression of the disease.

A key factor in the early identification of AD relates to its underlying neuropathology. Cellular studies of AD have consistently identified two hallmark features— extracellular deposits of beta-amyloid (*Aβ*) and intracellular neurofibrillary tangles composed of twisted tau filaments (Brion, 1998). The neural locations of these pathologies are not random. *Aβ* initially accumulates in medial prefrontal and medial parietal regions before progressing to temporal-lobe structures (Palmqvist et al., 2017). The tau filaments are first detected in temporal-lobe structures related to memory, including the entorhinal cortex, parahippocampal cortex, and hippocampus (Small et al., 2011). Importantly, these changes in the brain’s structure are often present before the appearance of clinical symptoms (Jack et al., 2013, 2019), thus highlighting the need for methods of early identification of at-risk individuals.

One intuitive method of early detection of Alzheimer’s susceptibility is genetic testing. Genetic researchers have identified numerous genes that increase an individual’s risk for developing AD (Breijyeh and Karaman, 2020). Specifically, Apolipoprotein (APOE)-*ε*4 has been found to be the strongest genetic risk factor for AD and is associated with increased levels of *Aβ* deposits and an earlier age of onset (M Di Battista et al., 2016). Structural neuroimaging studies found that these A*β* deposits affected the hippocampus; APOE-*ε*4 carriers exhibited smaller left hippocampal volumes than non-carriers of this gene (Wang et al., 2015). Additionally, a large body of work indicates that the phosphatidylinositol-binding clathrin assembly protein (PICALM) gene influences AD risk primarily by modulating the production, transport, and clearance of *Aβ* (Xu et al., 2015). Therefore, studying both APOE-*ε*4 and PICALM in genetically at-risk but otherwise healthy individuals could provide promising and novel avenues for understanding early AD diagnosis and for therapeutic development. For instance, a recent study (Morgen et al., 2014) found that the interaction between APOE-*ε*4 and PICALM rs3851179 adversely affects the cognitive performance (Trail Making Test A) and brain atrophy (prefrontal), respectively, in patients with early AD dementia. Thus, it is equally important to examine the functional impact of APOE-*ε*4 and PICALM rs3851179 on the brain to gain a more comprehensive understanding of their roles in early AD.

One neuroimaging tool that has been used to assess neural changes in AD is resting-state functional magnetic resonance imaging (rs-fMRI). This brain-imaging technique measures low-frequency fluctuations in the blood-oxygen-level-dependent (BOLD) signal while participants lie quietly without engaging in any task. Analysis of these BOLD fluctuations consistently shows a set of spatially distributed but temporally coherent restingstate networks (Raichle et al., 2001). Numerous restingstate networks have been identified. Prominent among these are the (1) Visual, (2) Sensorimotor, (3) Dorsal Attention, (4) Ventral Attention, (5) Salience (SN) Ventral Attention, (6) Limbic, (7) Frontoparietal, and (8) Defaultmode (DMN; see Raichle, 2015 (Raichle, 2015), for a review). In AD and mild cognitive impairment (MCI), disrupted connectivity has been most consistently observed in the DMN, arguably because some of its core hubs (i.e., posterior cingulate, hippocampus) are early sites of amyloid and tau pathology (Mintun et al., 2006). Studies finding alterations of rs-fMRI in the DMN (Allen et al., 2007; Greicius et al., 2004; Li et al., 2002) primarily have found decreased connectivity. A smaller body of work suggests that decreased connectivity in DMN may be accompanied by compensatory increase in connectivity in frontal executive regions (Agosta et al., 2012) or the SN (Zhou et al., 2010).

Importantly, the genetic variants that increase AD risk also modulate intrinsic brain networks before any clinical evidence of pathology. For example, one study observed that participants carrying an APOE-*ϵ*4 allele had clear indications of abnormalities in precuneus resting-state functional connectivity (FC) in the absence of any cognitive impairment, and in the absence of fibrillar cerebral (*Aβ*) deposits detectable by positron emission tomography imaging (Sheline et al., 2010). A recent study (Dzianok et al., 2025b) found that individuals carrying the APOE-*ϵ*4 allele exhibited reduced network strength in the posterior (temporo-occipital) portion of the middle temporal gyrus, compared with non-carriers, when independent components of the DMN were analyzed. Likewise, in a cohort of 283 young adults, PICALM G-allele carriers showed weaker connectivity compared with AA carriers in the hippocampal region (Zhang et al., 2015). However, although these studies provide important information about FC in individuals genetically at risk for AD, they estimate network connectivity by averaging patterns over the entire rs-fMRI scan (e.g., approximately 7-10 minutes). Recent advances in analysis techniques now enable researchers to track changes in FC throughout a restingstate scan (Hutchison et al., 2013; Preti et al., 2017). By capturing these brief connectivity states, they can directly link time-varying network changes to cognitive processes and behavior (Hutchison et al., 2013).

Dynamic resting-state functional connectivity (dFC) divides the BOLD time series into overlapping sliding windows (e.g., 30–60 s) and computes a separate connectivity matrix for each window (Hutchison et al., 2013; Preti et al., 2017). These windowed matrices are then clustered into a finite set of recurring connectivity states, each reflecting a distinct pattern of inter-regional coupling (Allen et al., 2012). From these states, we derive features such as dwell-time, the total time spent in each connectivity state. For example, Jones et al.(Jones et al., 2012) found a significant difference in dwell-time between AD and cognitively healthy groups. Another study found differences in dFC strength in the left precuneus, default mode network, and dorsal attention network among normal controls, aMCI patients, and AD patients (Zhao et al., 2022). Finally, linking dFC with genetic literature related to AD, Gong et al. (Gong et al., 2025) reported that APOE-*ϵ*4 carriers exhibit reduced dFC between the posterior cerebellar lobe and the middle temporal gyrus in AD. To date, no rsfMRI study has examined dFC changes associated with the PICALM gene; however, an electroencephalography (EEG) study (Ponomareva et al., 2020) found lower functional connectivity in PICALM GG carriers, suggesting early functional changes in the alpha frequency band.

In this study, we build upon this earlier dFC research to examine whether the dwell-time of resting-state networks varies across genotypes. Specifically, we use rsfMRI data from the Polish EEG, Alzheimer’s Risk-genes, Lifestyle and Neuroimaging (PEARL-Neuro) Database (Dzianok and Kublik, 2024) to compare healthy middle-aged individuals who had both the APOE-*ϵ*4 allele and the PICALM gene (i.e., the double-risk group), individuals with the APOE *ϵ*4 allele but not the PICALM gene, or individuals who did not have either AD-related allele. We specifically hypothesized that differences in dwell-time would be observed between the double-risk group and the no-risk group. These results would provide a novel biomarker for AD susceptibility and may provide insight into the neural mechanisms underlying the early symptoms of this disorder.

## Methods

### A. Participants

In this study, the PEARL-Neuro Database (Dzianok and Kublik, 2024), freely available online at the OpenNeuro database, is utilized. The complete PEARL dataset comprises 192 middle-aged individuals. However, only 69 subjects participated in the fMRI session. We further excluded 1 subject who carried the rare *ε*2/*ε*4 APOE haplotype, as these individuals possess both a risk allele (*ε*4) and a protective allele (*ε*2), creating an ambiguous genetic risk profile that does not conform to the predefined risk group classifications. This resulted in a final sample of 68 subjects for the current analysis. Our study specifically focuses on rs-fMRI data from these 68 healthy middle-aged subjects. In the previous study (Dzianok et al., 2025a), subjects were divided into three different categories based on their genetic risk factors for AD: (1) no-risk gene (APOE *ε*4/PICALM rs3851179 variants are absent, A-P- or N), (2) single-risk gene (APOE *ε*4 present/PICALM rs3851179 absent, A+P- ), (3) double-risk gene (APOE *ε*4/PICALM rs3851179 variants are both present, A+P+) to identify AD-like features in rs-EEG/fMRI data. The distribution of subjects in each group is shown in Table 1. For all subjects, the Edinburgh Handedness Inventory (EHI) test result confirmed right-handedness, and all subjects were generally healthy with no decline in cognitive task performance. This research received ethics approval from the University of Winnipeg Human Research Ethics Board.

**Table 1.** Description of the study participants by group.

| Group | Final Count | Ratio (%) | Age (Mean ± SD) | Sex Distribution |
| --- | --- | --- | --- | --- |
| N | 27 | 39.7 | 55.2 ± 2.9 | F: 15, M: 12 |
| A+P- | 23 | 33.8 | 55.7 ± 3.3 | F: 12, M: 11 |
| A+P+ | 18 | 26.5 | 55.7 ± 3.4 | F: 8, M: 10 |
| **Total** | **68** | **100.0** | **55.5 ± 3.1** | **F: 35, M: 33** |

### B. Scanning Parameters

The rs-fMRI and structural MRI (T1w and T2w) data from the PEARL database were recorded on a Siemens Prisma FIT 3 T scanner (Siemens Medical Systems, Erlangen, Germany) using a gradient-echo EPI sequence with repetition time (TR) = 0.8 s, echo time (TE) = 38 ms, slice thickness = 2 mm, and voxel size = 2 ×2 ×2 mm^3^. Images were taken with two-phase encoding directions: Anterior-to-Posterior (AP) and Posterior-to-Anterior (PA) for approximately 7.5 minutes each.

### C. fMRI Preprocessing

First, since the fMRI data were recorded using two-phase encoding directions (AP and PA), we corrected the geometric distortions that occur in fMRI scans by applying FSL topup correction (Andersson et al., 2003). These distortions are usually caused by static magnetic field inhomogeneities, leading to pixel shifts, particularly in the phase encode direction (Jezzard and Balaban, 1995). The FSL top-up correction was conducted by taking a mean of the whole AP and PA scans (volume dimensions 108 ×108 ×72 voxels), and then estimating distortions in the AP and PA rs-fMRI scans for each subject. After the distortions were found, we applied a top-up method to correct these distortions from both scans for each subject.

Subsequently, we used a well-established pipeline, fMRIprep (Esteban et al., 2019, 2020), to further enhance our data preprocessing. The undistorted data from AP and PA scans underwent the following steps using fMRIprep: a reference volume was generated and used to estimate and correct head motion (via FSL mcflirt); each scan was then co-registered to the subject’s T1w image using FreeSurfer’s boundary-based registration and FSL flirt (6 degrees of freedom), and all spatial transforms (motion, distortion correction, anatomical alignment and normalization to Montreal Neurological Institute (MNI) space) were composed into a single interpolation step. The coregistration template that we used was the MNI152Nonlinear2009cAsym template (Fonov et al., 2011, 2009). The final volume dimensions after fMRIprep, both AP and PA scans, were 97 ×115× 97 ×560 voxels, where the first three dimensions were spatial (voxels) and the fourth dimension (*t* = 560) was the number of time points sampled with TR = 0.8s.

Alongside the preprocessed images, fMRIPrep generates a set of confound regressors for each functional run, which were not removed in prior steps. From those regressors, six head motion parameters, translations (x, y, z) and three rotations (pitch, roll, yaw), were selected and subsequently regressed out from the BOLD timeseries data to mitigate the effects of subject motion. In addition, we regressed out physiological-noise components using aCompCor signals derived from subject-specific white-matter and CSF masks (provided by fMRIPrep) Behzadi et al. (2007). These regressors were removed from each ROI time series before computing dFC. Because respiration and cardiac recordings were not available, RETROICOR could not be applied; therefore, aCompCor was used as the primary physiological-noise mitigation strategy. This choice is further motivated by the fact that aCompCor does not require external physiological monitoring, which is advantageous when such recordings are unavailable Behzadi et al. (2007).

### D. Correlation Computation and Clustering

After the AP and PA scans were cleaned, we extracted correlation matrices across seven brain networks: (1) Visual, (2) Somatomotor, (3) Dorsal Attention, (4) Salience/Ventral Attention, (5) Limbic, (6) Control, and (7) DMN. These networks are used in many fMRI studies, and they involve a wide range of brain structures, some of which are closely associated with AD. For instance, the hippocampus, part of several networks and a sub-region of the DMN, often shows weaker FC and structural loss in AD (Andrews-Hanna et al., 2010; Zhou et al., 2008; O’Callaghan et al., 2019; Lee et al., 2020). Similar AD-related changes have been found in the other six networks, motivating our decision to focus on this set of seven large-scale networks for subsequent analyses.

We used the 100-parcel, 7-network Schaefer atlas, which is aligned with the MNI-152 Nonlinear 2009cAsym template employed in preprocessing, and subdivides each large-scale network into smaller cortical subregions (e.g., medial prefrontal cortex, precuneus). This atlas provides 100 non-overlapping parcels that cover the whole cortex. Using these 100 parcels, we computed sliding-window correlations. The sliding window moved across the entire duration of each scan using a fixed 60 s window (60/0.8 = 75 volumes) with 75% overlap. With 75% overlap, the step size is 15 s, which corresponds to 18.75 volumes; therefore, we used a 19-volume step (approximately 75% overlap). A 75% overlap was chosen to better capture short or boundary-adjacent fluctuations in FC and to reduce variance in our correlation estimates. Consequently, for each subject, the resulting correlation matrices across both scans had dimensions 100 ×100 ×52, where 52 was the total number of windows across both scans (26 per scan). This was computed as 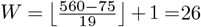, where 560 is the number of volumes per scan, 75 is the window size in volumes, and 19 is the step size in volumes. For each window *w* = 1,…, *W*, the correlations across all parcellated regions were computed by first calculating the pairwise covariance and variances in window *w*:

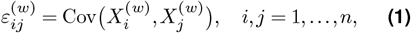

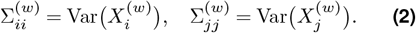

Then the Pearson correlation for each pair (*i, j*) is

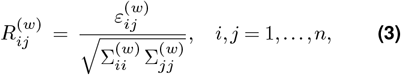

where 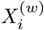 is the time series of parcel *i* in window *w*, 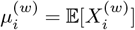 its mean, and 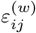 the covariance between parcels *i* and *j* in that window.

Finally, since correlation FC does not contain any directional information and produces a symmetric (mirror) square matrix, we only used the lower-triangular entries and used these feature vectors as input to K– means clustering with *K* clusters. Rather than selecting a single fixed value of *K*, we explored a range of cluster numbers. The lower bound of this range was informed by the elbow method (Figure 1), which identifies a point of diminishing returns in within-cluster variance reduction. The upper bound was determined empirically by examining the distribution of subjects across clusters: beyond a certain value of *K*, some clusters contained very few windows from only a handful of subjects, reducing statistical power and interpretability. The shaded region in Figure 1 reflects this valid *K*-range, bounded by values where cluster formation remained both stable and statistically meaningful for group-level analysis. To visualize the clusters, we applied principal component analysis (PCA) to reduce the 4950 (100*99/2) connectivity pairs to two dimensions and plotted each sample by its cluster assignment to visualize cluster formation.

**Figure 1.**
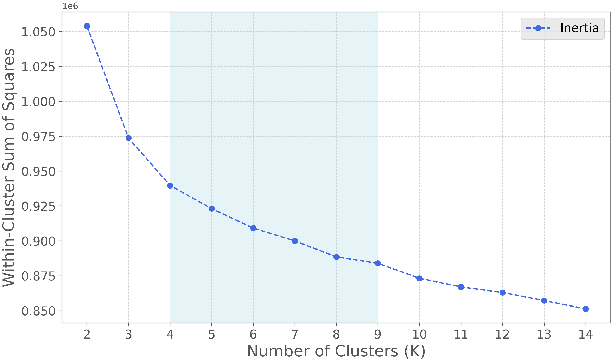
Elbow method for determining the optimal number of clusters (*k*) using K-means clustering. The y-axis represents the within-cluster sum of squares (WCSS), or inertia, while the x-axis indicates the cluster count. The shaded region (clusters 4–9) denotes the optimal range selected for subsequent statistical analysis and machine learning classification. Abbreviations: *k*: cluster count and WCSS: within-cluster sum of squares.

### E. Feature Extraction and Statistical Methods

First, we applied K-means clustering to the slidingwindow dFC observations to assign each window to one of *K* connectivity states (clusters). Here, each slidingwindow dFC observation corresponds to a window-level FC matrix (i.e., a short-time “snapshot” of inter-regional connectivity). K-means groups windows with similar FC patterns, and each resulting cluster represents a recurring FC pattern. Next, to estimate the total duration each subject spent in a specific state while accounting for the overlap between adjacent windows, we computed the dwell-time (DT_*k*_) using the temporal stride. The dwell-time for cluster *k* was defined as:

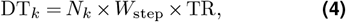

where *N*_*k*_ denotes the number of windows assigned to cluster *k*, and *W*_step_ represents the sliding-window step size in volumes. By multiplying the step size in volumes by the repetition time (TR = 0.8 s*/*volume), we obtain the effective temporal duration contributed by each window (the stride). Specifically, we used a window length of 60 s (75 volumes) with a 75% overlap, resulting in a step size of *W*_step_ = 19 volumes. This corresponds to a temporal stride of approximately 15 s (i.e., 19 × 0.8 ≈15.2 s). Thus, DT_*k*_ represents the cumulative time (in seconds) that a subject’s brain activity was classified into state *k*.

We repeated the clustering and subsequent dwell-time analysis for multiple choices of *K* to evaluate whether findings were consistent across different numbers of states. For each value of K, we tested whether dwell-time for each cluster differed across genetic risk groups (N, A+P- and A+P+) using an adaptive statistical approach. For each cluster, we first assessed the assumptions required for parametric testing by applying the Shapiro-Wilk test for normality within each group and Levene’s test for homogeneity of variance across groups. When both normality (p *>* 0.05) and variance homogeneity (p *>* 0.05) assumptions were satisfied, we used parametric tests: t-test for two-group comparisons (for machine learning) or ANOVA for multi-group comparisons (for initial statistical analysis). When these assumptions were violated, we employed non-parametric alternatives: the Mann-Whitney U test for two groups or the Kruskal-Wallis test for multiple groups. To control the family–wise error across the K clusters, the resulting p–values were corrected using the Benjamini— Hochberg false discovery rate (FDR) procedure. Clusters with FDR-corrected (p *<* 0.05) were considered significant.

### F. Hubness Analysis via Statistical Testing

After the K-means clustering, we selected clusters producing significant or lowest p-values between-group differences (p *<* 0.05) in dwell-time and extracted their 1D correlation vectors, then reconstructed them into symmetric 100 ×100 connectivity matrices for each sample within those clusters. We then computed the global mean connectivity matrix by averaging all connectivity matrices (alternatively, by using the cluster centroid) across all subjects and groups within each significant cluster, under the assumption that clustered windows reflect a common network configuration that transcends group boundaries while still showing differential dwell-time patterns. This global approach allows us to measure the underlying connectivity that defines each significant cluster, providing insight into the network topology that groups differentially occupied in time. As the dwell-time feature represents how much time a subject spent in a given cluster, we chose to use hubness analysis on this global connectivity pattern to identify dominant “sink” regions that measure each network configuration. This approach helps link temporal dynamics (i.e., dwell-time differences between groups) with spatial topological features (i.e., globally consistent hub regions), providing a unified interpretation of how genotype-specific temporal patterns emerge within shared connectivity configurations.

To identify the top 10 hub regions within each significant cluster, we established a threshold by computing the median plus two standard deviations of this global mean connectivity matrix, which corresponds to approximately the top 2—3% of connectivity values. We then binarized the global connectivity matrix using this threshold and computed hubness proportions for each region by summing its corresponding row in the binary matrix and dividing by the total number of regions minus one (99). The ten regions with the highest hubness proportions from each significant (lowest p-value) cluster were aggregated across all K values and multiple random seeds to identify the most consistently appearing hub regions.

### G. Machine learning

As a complementary analysis, we applied a simple machine learning classifier to test whether dwell-time features could generalize to unseen data. Specifically, we used logistic regression to predict genotype groups based on dFC dwell-time profiles. This analysis helped evaluate the potential of these functional markers in distinguishing at-risk individuals, beyond group-level statistics. Figure 2 illustrates the comprehensive workflow for genotype prediction using dwell-time features. We framed the genotype prediction task as a set of binary classifications comparing groups N, A+P-, and A+P+ as N vs A+P-, N vs A+P+, A+P- vs A+P+, and N vs (A+P- and A+P+ combined). The overall pipeline consists of several sequential steps:

**Figure 2.**
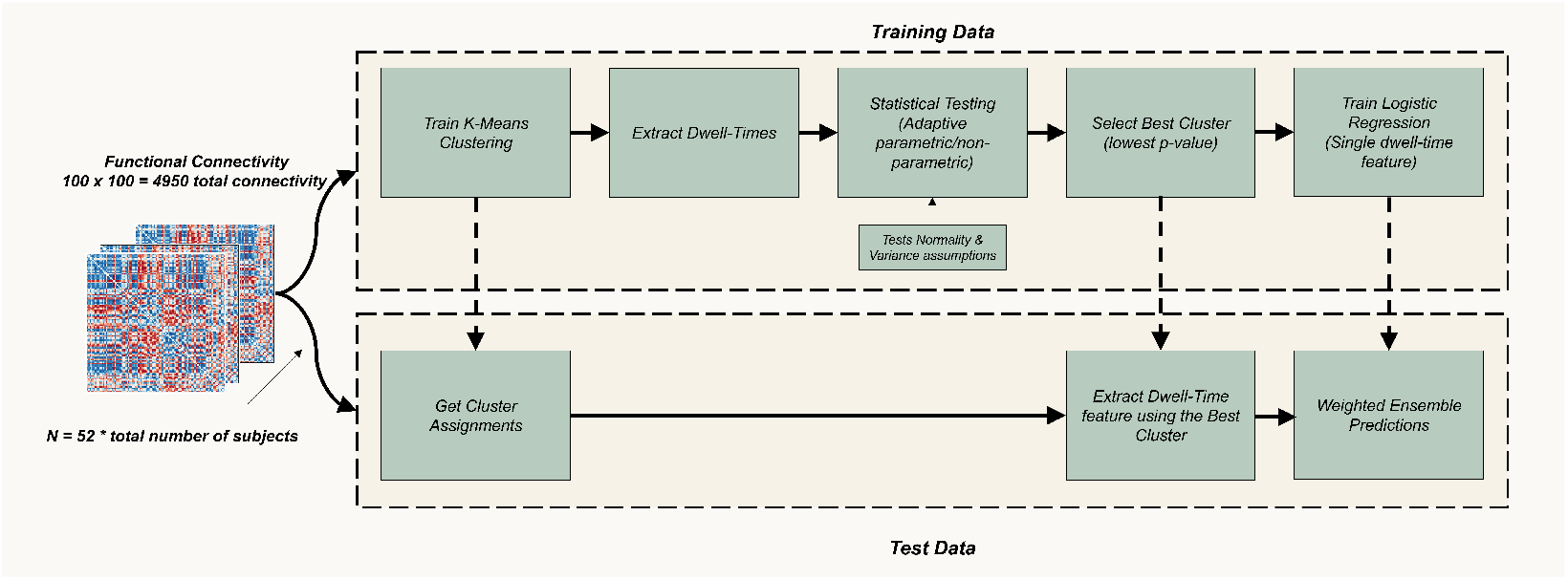
Proposed dFC classification pipeline. Flowchart of the machine learning approach for distinguishing genetic risk groups. The pipeline utilizes K-means clustering to define connectivity states, followed by adaptive statistical testing (Shapiro-Wilk for normality and Levene’s for variance) to select the most discriminative cluster (*k* = 4 9) based on dwell-time features. Final predictions are generated via a weighted ensemble of logistic regression models. Abbreviations: dFC: dynamic functional connectivity and *N* : number of dFC windows per subject (*N* = 52).

#### G.1 Data Preprocessing and Cross-Validation Setup

First, we ensured that the FC matrices (100 ×100) were flattened into 1D vectors containing 4,950 unique connectivity features, which served as the required input format for k-means clustering. Subjects were then partitioned into training and testing folds using stratified 10-fold cross-validation.

#### G.2 Training Phase

Within each training fold, k-means clustering was performed exclusively on training data to identify K connectivity clusters (where K ranges from 4 to 9, see Figure 1). Based on these cluster assignments, we computed each subject’s dwell-time, the total time spent in each connectivity cluster. To select the most discriminative cluster, we employed adaptive statistical testing that first evaluates normality (ShapiroWilk test) and variance homogeneity (Levene’s test) assumptions. Depending on whether these assumptions were satisfied, we applied either parametric tests (t-test) or non-parametric alternatives (Mann-Whitney U test). The cluster with the lowest p-value was selected, and the dwell-time feature was extracted. Finally, the single dwell-time feature was used to train a logistic regression model. Therefore, in one run, six logistic regression models trained considering the K = 4 to 9 range were used.

#### G.3 Test Phase

Crucially, the same k-means model trained on the training data was applied to assign test subjects to clusters, ensuring no data leakage occurs since test data never influences cluster construction. Dwell-time features were extracted from test subjects using the previously identified optimal cluster, and predictions were generated using the corresponding trained logistic regression model.

#### G.4 Ensemble Method

This process was repeated across K = 4 to 9 clusters and five random seeds for stability, resulting in six logistic regression models per seed. To combine predictions across different K values for one seed run, we implement a p-value weighted ensemble method where models with lower p-values received higher weights in the final prediction. The weights were computed as follows:

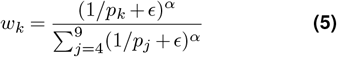

Where *p*_*k*_ was the p-value for cluster *K, p*_*j*_ was the p-value for cluster j in the denominator sum, where j ranges from 4 to 9, *ε* = 10−10 was a smoothing parameter to prevent division by zero, and *α* = 2 was the power parameter that emphasizes models with lower p-values. Finally, the prediction was computed as follows:

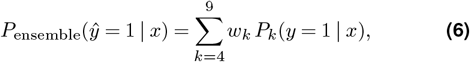

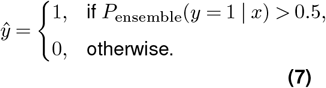

where *P*_*k*_(*ŷ* = 1 |*x*) was the predicted probability from the logistic regression model trained using *K* clusters, and *ŷ*_ensemble_ was the final binary prediction. This weighted-ensemble approach prioritizes models that demonstrate stronger statistical evidence for group differences.

### H. Performance Evaluation and Model Interpretability

The performance was evaluated using accuracy, precision, recall, F1-macro and Area under the Receiver Operating Characteristic Curve (AUC-ROC). For each seed, we averaged each metric across the 10 folds; the final reported score for each metric is the mean ± standard deviation across the five seeds. All preprocessing (including feature standardization) and feature selection were performed using training data only. For reproducibility, K-means initialization and fold generation were seeded (with different seeds) for each run.

To show the interpretation of our logistic regression models, we extracted coefficients from each fold and back-transformed them from scaled values to their original units of seconds. This transformation enabled the interpretation of how dwell-time in statistically significant connectivity clusters correlates with genetic risk group membership. We computed effect sizes at the 60second time scale, corresponding to our sliding window length. The resulting coefficient indicated the change in log-odds per additional 60 seconds of dwell-time.

#### H.1 Permutation Test

To assess the statistical significance of our classification results and ensure they were not due to chance, we performed a permutation test for each classification task. This procedure tests the null hypothesis that there is no true association between dwell-time features and genotype labels. To test this, for each task, the entire classification pipeline was first run with the correct subject-label assignments to calculate the observed AUC. Then, the procedure was repeated 1,000 times. In each repetition, the genotype labels were randomly shuffled across subjects, thereby breaking any true relationship between the features and the labels, while preserving the data shape of dwell-time features and the imbalance between groups. This process generated a null distribution of AUC scores, representing the range of performance expected purely by chance. The empirical p-values are calculated as follows:

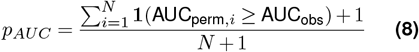

This is an empirical permutation testing approach where the p-value represents the probability of observing a test statistic (AUC) as high or higher than the observed value under the null hypothesis of no group differences.

## Results

### I. Statistical Analysis

Our statistical analysis was focused on dwell-time features derived from k-means clustering using dFC windows, which were evaluated across a range of k values (k = 4 to 9). This analysis consistently revealed two distinct clusters that demonstrated opposing patterns of dwell-time between subject groups (see Figure 3). For instance, at k=5, we observed that cluster 3 was characterized by significantly longer dwell-times for the N group compared to the A+P- and A+P+ group, whereas cluster 0 was characterized by marginally significant shorter dwell-times for the N group compared to the A+P-group. This pattern of two opposing clusters, one with increased dwell-time for the N group and another with decreased dwell-time, was a consistent trend across most k-values and random seeds. The large distance between the centroids of these two clusters suggests they represent fundamentally different functional connectivity patterns rather than minor variations of a single cluster. It is important to note that while these group differences were prominent, they did not survive the primary statistical correction for multiple comparisons across all clusters, and only some survived posthoc p-value correction.

**Figure 3.**
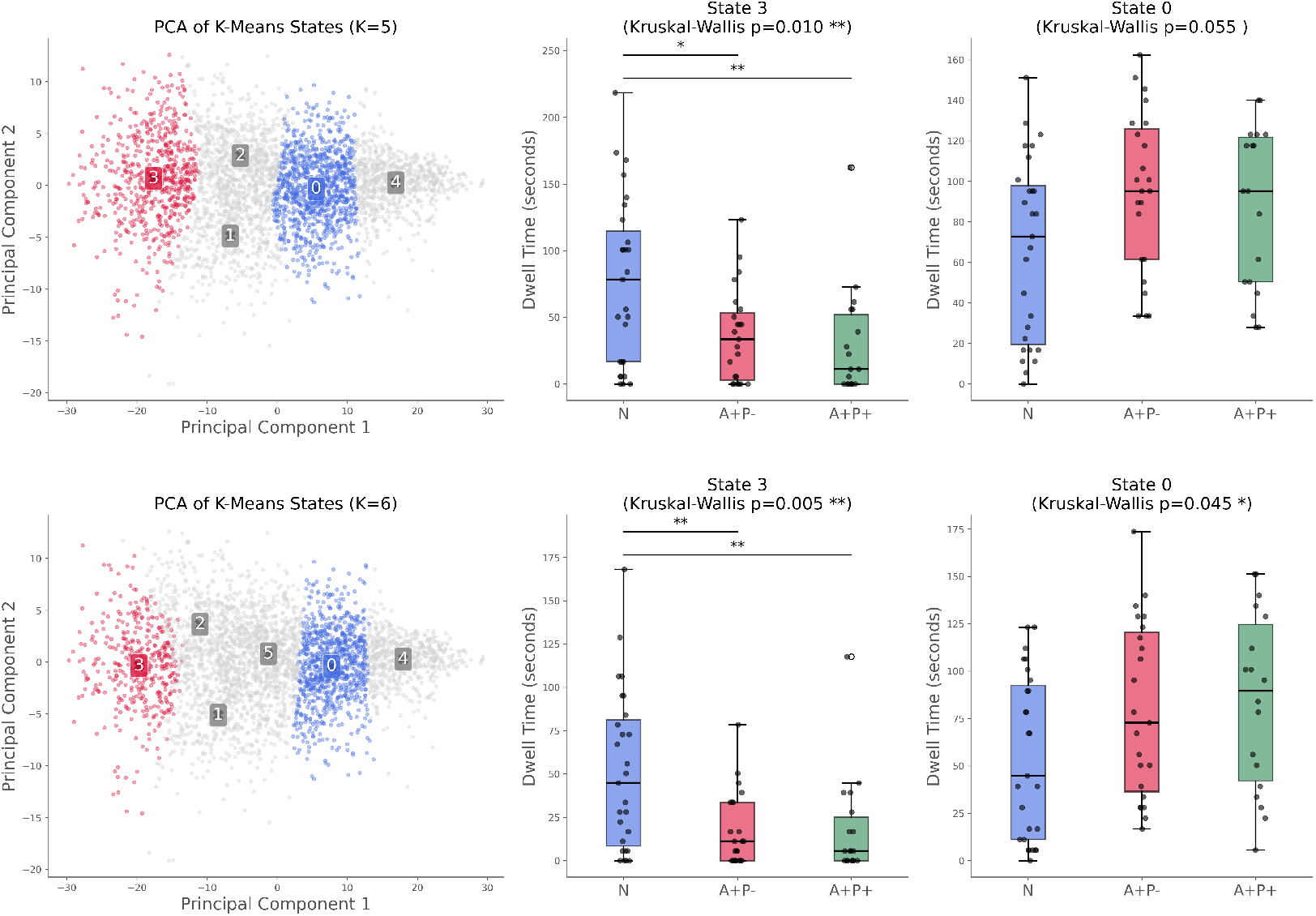
Discriminative dFC clusters and group dwell-time comparisons for *k* = 5 and *k* = 6. Left panels show Principal Component Analysis (PCA) projections of FC windows. Clusters are colored based on statistical significance: red indicates significantly higher dwell-time for the Not-at-risk (N) group, blue indicates higher dwell-time for at-risk groups (A+P-/A+P+), and gray represents non-significant clusters. Right panels display box plots comparing total dwell-time (seconds) between groups. Statistical significance was determined using Kruskal-Wallis tests, with asterisks denoting significance levels (* : *p <* 0.05, ** : *p <* 0.01). Abbreviations: PCA: Principal Component Analysis, FC: functional connectivity, N: not-at-risk group, A+P-: APOE+ PICALM-, and A+P+: APOE+ PICALM+.

### J. Hubness Analysis

To understand which brain regions best separated the groups, we looked for regions that repeatedly showed up as highly connected “hub” regions within the important clusters. Tables 2 and 3 list the top 10 hub regions for: (1) clusters where the N group spent more time (higher dwell-time), and (2) clusters where the N group spent less time (lower dwell-time). Results were combined across all *k* values and five random seeds. In the clusters where the N group had higher dwell-time than A+P- and A+P+, the hubs were mainly on the left hemisphere of the brain, 9 out of the top 10 regions. In the clusters showing the opposite pattern (where the N group had lower dwell-time than the other groups), the hub regions were more evenly split between the left and right hemispheres. We also found that hub regions were more consistent in the clusters where the N group had higher dwell-time. This means the same regions appeared more often in the top 10 list, which suggests this connectivity pattern may occur more reliably across subjects and random seeds. Figures 4(a) and 4(b) show where these hub regions are located. Across both cluster types, regions from the Dorsal Attention network and the Salience/Ventral Attention network were common among the top hubs. In the clusters where the N group had higher dwell-time, Visual Cortex region 1 showed the strongest hub behaviour (average of 9.6 connections), and Dorsal Attention (posterior) region 3 was also strongly connected (average of 8.0 connections). In contrast, in the clusters where A+P- and A+P+ had higher dwell-time than the N group, hub regions were mostly from attention-related networks: 8 out of the top 10 hubs belonged to attention networks with weaker hub ratios.

**Table 2.**
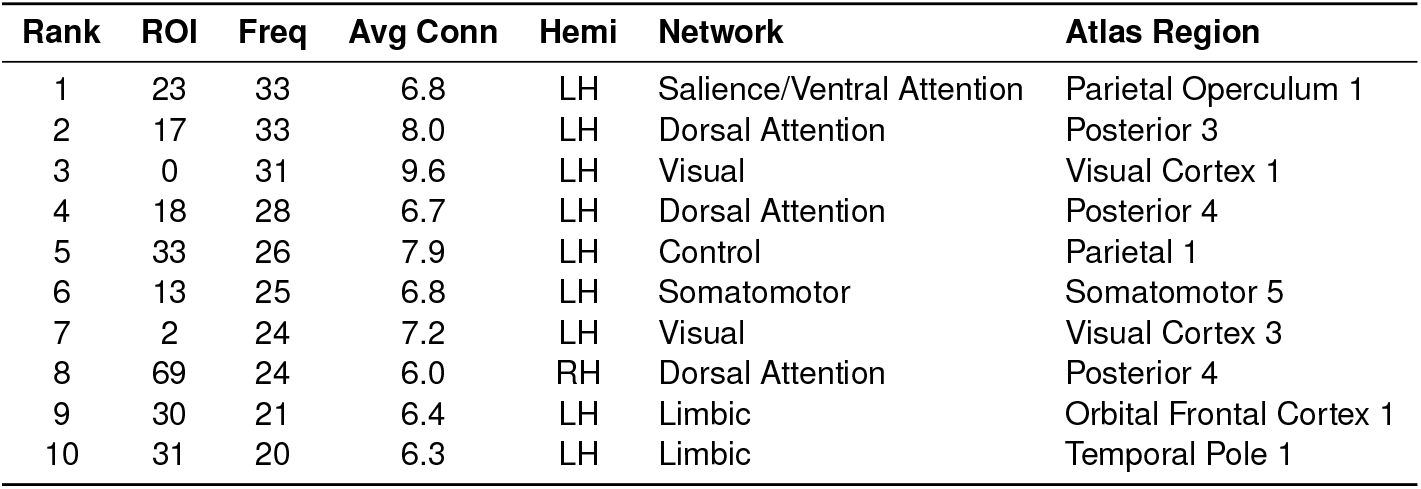
Top 10 Most Frequent Hub Regions (N Group Higher dwell-time clusters). ROI indices and region names correspond to the Schaefer (2018) 100-parcel atlas. The *Freq* column shows frequency across all runs (across *K* values and seeds), and the *Avg Conn* column indicates average connectivity strength for that region.

**Table 3.**
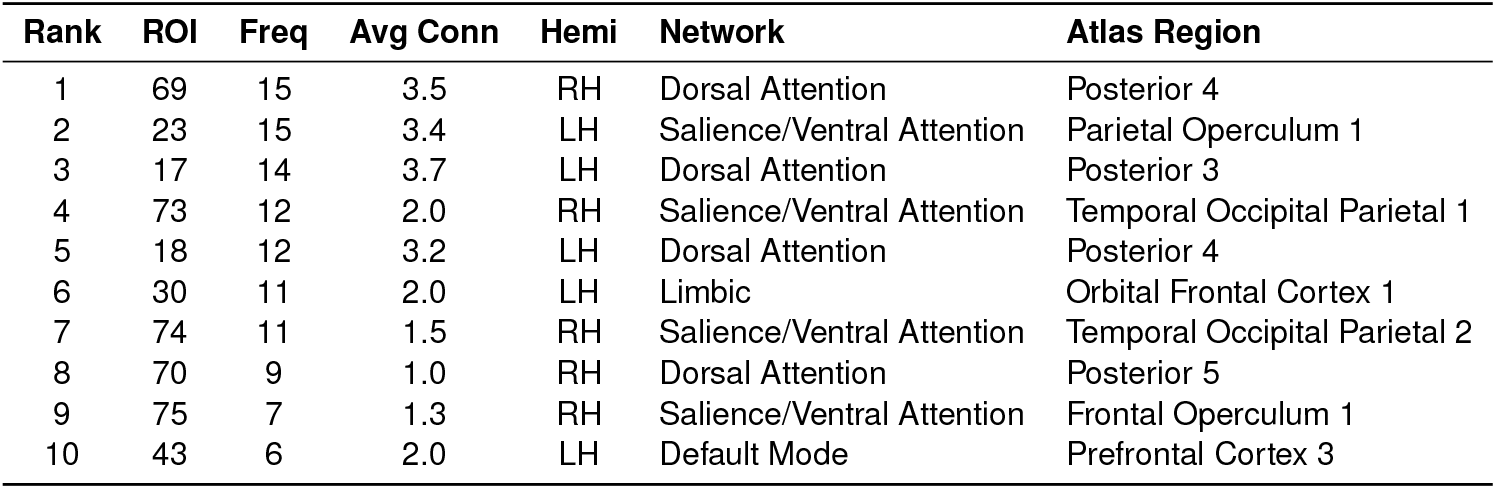
Top 10 Most Frequent Hub Regions (N Group Lower dwell-time clusters). ROI indices and region names correspond to the Schaefer (2018) 100-parcel atlas. The *Freq* column shows frequency across all runs (across *K* values and seeds), and the *Avg Conn* column indicates average connectivity strength for that region.

| Rank | ROI | Freq | Avg Conn | Hemi | Network | Atlas Region |
| --- | --- | --- | --- | --- | --- | --- |
| 1 | 69 | 15 | 3.5 | RH | Dorsal Attention | Posterior 4 |
| 2 | 23 | 15 | 3.4 | LH | Saliency/Ventral Attention | Parietal Operculum 1 |
| 3 | 17 | 14 | 3.7 | LH | Dorsal Attention | Posterior 3 |
| 4 | 73 | 12 | 2.0 | RH | Saliency/Ventral Attention | Temporal Occipital Parietal 1 |
| 5 | 18 | 12 | 3.2 | LH | Dorsal Attention | Posterior 4 |
| 6 | 30 | 11 | 2.0 | LH | Limbic | Orbital Frontal Cortex 1 |
| 7 | 74 | 11 | 1.5 | RH | Saliency/Ventral Attention | Temporal Occipital Parietal 2 |
| 8 | 70 | 9 | 1.0 | RH | Dorsal Attention | Posterior 5 |
| 9 | 75 | 7 | 1.3 | RH | Saliency/Ventral Attention | Frontal Operculum 1 |
| 10 | 43 | 6 | 2.0 | LH | Default Mode | Prefrontal Cortex 3 |

**Figure 4.**
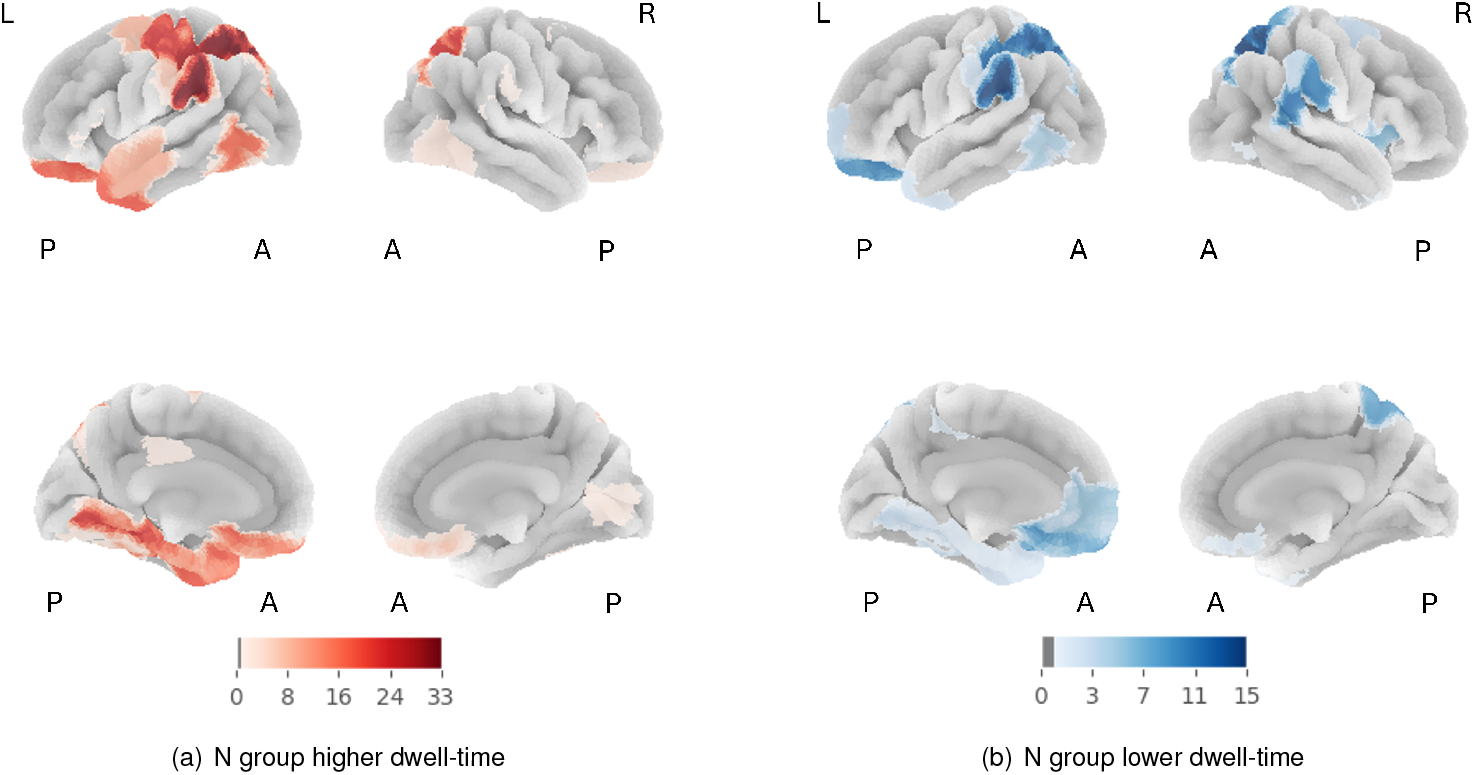
Comparative hub frequency maps for dFC clusters. Left panel (a): hub regions for clusters where the not-at-risk group (N) shows higher dwell-time; Right panel (b): hub regions for clusters where the not-at-risk group (N) shows lower dwell-time. Colour intensity indicates the frequency of hub detection across multiple *K* values, defined as the number of *K* values (from *K* = 4 to *K* = 9) for which the region was identified as a significant hub (*p <* 0.05). To assess stability, the full clustering and hub-identification procedure was repeated five times with different random seeds, and the resulting hub patterns were consistent across runs. “L” and “R” indicate left and right hemispheres, and “A” and “P” indicate anterior and posterior. The top row shows lateral views, and the bottom row shows medial views.

To assess whether the hubness pattern differed between the two cluster types (i.e., clusters where the N group showed higher dwell-time vs. clusters where the N group showed lower dwell-time relative to A+P- and A+P+), we compared the distributions of hub frequencies across ROIs. Across the aggregated top-10 hub rankings (all *k* values and five random seeds), 35 unique ROIs were observed. The overall hub-frequency distribution differed significantly between the two cluster types (*χ*^2^(34) = 176.08, *p <* 0.001), indicating that they are characterized by distinct hub configurations. Given the small per-ROI counts and the aggregation across multiple *k*/seed runs, we interpret this result primarily as evidence of a global shift in hub organization rather than attributing the effect to specific regions. For completeness, we examined standardized residuals from the contingency table as a post-hoc check (threshold ± 1.96, *p <* 0.05), which suggested that only a small subset of ROIs showed disproportionate hub frequencies (i.e., how often a region appeared in the top-10 list) between the two cluster types. Overall, the difference appears distributed across many ROIs, supporting the view that these two cluster types reflect fundamentally different connectivity states.

Importantly, these statistical dwell-time findings show associative patterns rather than causal relationships. Although we cannot establish directionality between brain connectivity and genotype, the systematic differences in dwell-time features provide discriminative ability that can be exploited for genotype prediction and further interpretation. To test this, we used logistic regression, which provides both predictive performance and interpretability. The sign and magnitude of each coefficient, and the corresponding odds ratio, describe how the odds of a genotype change for a one-unit (60 s) increase in a feature, providing a clear effect-size interpretation. In the subsequent machine-learning analysis, we therefore evaluate predictive performance and report coefficients and odds ratios to identify robustness and interpretability of dwell-time features.

### K. Machine Learning

#### K.1 Classification Performance

We evaluated whether dwell-time features derived from dFC can predict APOE *ε*4/PICALM genotype categories by training binary logistic regression models with subject-stratified 10-fold cross-validation repeated five times. Overall performance across tasks was modest but above chance for comparisons involving the non-carrier group (N; Table 4). The strongest discrimination was observed for N vs. (A+P-& A+P+) with an accuracy of 0.682 ± 0.025 and an AUC-ROC of 0.696 ± 0.022. Among the pairwise comparisons, N vs. A+P+ showed the highest AUC-ROC (0.680 ± 0.024) and competitive accuracy (0.618 ± 0.029), while N vs. A+P-was weaker but still above chance (accuracy 0.596 ± 0.048; AUC-ROC 0.636 ± 0.021). In contrast, the A+P-vs. A+P+ task remained near chance-level (accuracy 0.507 ± 0.105; AUC-ROC 0.479 ± 0.082), indicating limited separability between the single- and double-risk genotype groups using dwell-time features alone.

**Table 4.** Model performance (mean *±* std) for binary classification tasks. Results computed over subject-stratified 10-fold CV repeated 5 times.

| Metric | N vs A+P- | N vs A+P+ | A+P- vs A+P+ | N vs (A+P- & A+P+) |
| --- | --- | --- | --- | --- |
| Accuracy | 0.596 ± 0.048 | 0.618 ± 0.029 | 0.507 ± 0.105 | **0.682 ± 0.025** |
| Precision | 0.601 ± 0.046 | 0.627 ± 0.028 | 0.502 ± 0.101 | **0.669 ± 0.025** |
| Recall | 0.600 ± 0.047 | 0.631 ± 0.029 | 0.499 ± 0.099 | **0.670 ± 0.024** |
| F1 (macro) | 0.595 ± 0.048 | 0.616 ± 0.029 | 0.497 ± 0.099 | **0.669 ± 0.025** |
| AUC-ROC | 0.636 ± 0.021 | 0.680 ± 0.024 | 0.479 ± 0.082 | **0.696 ± 0.022** |

#### K.2 Coefficient Orientation

To interpret which dwell-time patterns contributed most to classification, we summarized the logistic regression coefficients by the *direction* of the dwell-time difference (Table 5). Coefficients are reported as the change in log-odds per 60 seconds of additional dwell-time, averaged across five random seeds.

**Table 5.** Logistic regression coefficients by dwell time direction. Coefficients represent log-odds change per 60 seconds of additional dwell time. Values are mean ± SD across 5 seeds. All p-values *<* 0.05 after FDR correction.

| Direction | Task | n | $\beta$ (per 60s) | OR | p (FDR) |
| --- | --- | --- | --- | --- | --- |
| N/A+P- higher<br>(negative $\beta$) | N vs. A+P- | 218 | -0.673 ± 0.048 | 0.533 | $< 0.001$ |
| | N vs. A+P+ | 286 | -0.533 ± 0.027 | 0.605 | $< 0.001$ |
| | A+P- vs. A+P+ | 195 | -0.333 ± 0.159 | 0.763 | 0.009 |
| | N vs. (A+P- & A+P+) | 281 | -0.547 ± 0.025 | 0.593 | $< 0.001$ |
| A+P/A+P+ higher<br>(positive $\beta$) | N vs. A+P- | 82 | 0.454 ± 0.053 | 1.613 | $< 0.001$ |
| | N vs. A+P+ | 14 | 0.298 ± 0.069 | 1.355 | $< 0.001$ |
| | A+P- vs. A+P+ | 105 | 0.664 ± 0.151 | 3.746 | $< 0.001$ |
| | N vs. (A+P- & A+P+) | 19 | 0.359 ± 0.052 | 1.434 | $< 0.001$ |

In *Direction 1* (clusters where the N or A+P-group showed higher dwell-time), coefficients were negative and odds ratios (OR) were *<* 1, meaning that spending more time in these clusters was associated with *lower* odds of belonging to the comparison risk group. This pattern was consistent and significant across tasks: N vs. A+P-(*β* = ™0.673 ± 0.048, OR= 0.533), N vs. A+P+ (*β* = ™0.533 ± 0.027, OR= 0.605), and N vs. (A+P-& A+P+) (*β* = ™0.547 ± 0.025, OR= 0.593; all *p*_FDR_ *<* 0.001). The A+P-vs. A+P+ task also reached significance in this direction (*β* = ™0.333 ± 0.159, OR= 0.763, *p*_FDR_ = 0.009), indicating that some cluster configurations favored A+P-over A+P+.

In *Direction 2* (clusters where A+P-/A+P+ showed higher dwell-time than N, or A+P+ exceeded A+P-), coefficients were positive and OR were *>* 1, meaning that spending more time in these clusters was associated with *higher* odds of belonging to the risk group. This direction was also significant across tasks, including N vs. A+P-(*β* = 0.454 ±0.053, OR= 1.613), N vs. A+P+ (*β* = 0.298 ±0.069, OR= 1.355), A+P-vs. A+P+ (*β* = 0.664 ±0.151, OR= 3.746), and N vs. (A+P-& A+P+) (*β* = 0.359 ±0.052, OR= 1.434; all *p*_FDR_ *<* 0.001). Overall, these results show a clear and interpretable pattern: clusters where the N group spent more time were associated with reduced odds of risk-group membership, whereas clusters where the risk groups spent more time were associated with increased odds of risk-group membership.

#### K.3 Interpreting Odds Ratios

All odds ratios in Table 5 are reported per 60-second increase in dwell-time. OR *<* 1 indicates that more dwell-time in that cluster type is associated with lower odds of being in the risk group (i.e., relatively higher odds of being in N for contrasts involving N). For example, in *Direction 1*, OR= 0.533 for N vs. A+P-corresponds to a ≈ 46.7% reduction in the odds of belonging to A+P-per additional 60 seconds of dwell-time, and OR= 0.593 for N vs. (A+P-& A+P+) corresponds to a ≈40.7% reduction in odds of belonging to the combined risk group. Conversely, OR *>* 1 indicates higher odds of belonging to the risk group as dwell-time increases. For example, in *Direction 2*, OR= 1.613 for N vs. A+P-corresponds to a ≈61.3% increase in odds of belonging to A+P-per additional 60 seconds of dwell-time, and OR= 1.434 for N vs. (A+P-& A+P+) corresponds to a ≈ 43.4% increase in odds of belonging to the combined risk group. Together, the direction-specific results show that dwell-time carries consistent information about genotype group differences, and that the sign of the coefficient provides an intuitive interpretation of whether a cluster type is more characteristic of the N group or the risk groups.

#### K.4 Permutation Test

To assess whether dFC dwell-time features could reliably classify subject groups beyond chance, we evaluated statistical significance using a non-parametric permutation testing framework with 1000 permutations (genotype labels shuffled). The observed AUC values and corresponding empirical p-values are summarized in Table 6. Two comparisons reached statistical significance: N vs. A+P+ (AUC = 0.680, *p* = 0.034) and N vs. (A+P-& A+P+) (AUC = 0.696, *p* = 0.005), indicating that these classification results are unlikely to arise from random label assignments. In contrast, N vs. A+P-showed a trend-level effect (AUC = 0.636, *p* = 0.067) but did not meet the conventional threshold (*p <* 0.05), and A+P-vs. A+P+ was consistent with chance performance (AUC = 0.479, *p* = 0.486). Overall, the permutation test supports that dwell-time features provide reliable discriminatory signal primarily for contrasts involving the non-carrier group (N), particularly when comparing N againsts A+P+ or against the pooled risk groups.

**Table 6.** Summary of Classification Performance and Permutation Test Results.

| Comparison | Observed AUC ↑ | Empirical p-value ↓ |
| --- | --- | --- |
| N vs. A+P- | 0.636 | 0.067 |
| N vs. A+P+ | **0.680** | **0.034\*** |
| A+P- vs. A+P+ | 0.479 | 0.486 |
| N vs. (A+P- & A+P+) | **0.696** | **0.005\*\*** |

## Discussion

Our analysis of dwell-time profiles revealed robust group differences across varying numbers of clusters (*K* = 4 to 9) and multiple random seeds. Rather than relying on a single fixed cluster index, we observed that the resulting connectivity states consistently aligned into two distinct categories: (1) states where the not-at-risk (N) group exhibited significantly longer dwell-times than the at-risk groups (*A* + *P*™ and *A* + *P* +), and (2) states showing the opposite pattern, where at-risk groups spent more time than the N group. This recurrence across different model orders (*K*) confirms that these profiles represent stable biological signals rather than artifacts of a specific parameter choice. Furthermore, the clear separation of cluster centroids (see Figure 3) supports their interpretation as functionally distinct connectivity patterns. Importantly, we evaluated the full range of *K* values empirically, without selecting specific configurations, ensuring that our results are unbiased and reproducible.

For the interpretation of these clusters, which showed recurring significant cluster patterns, we identified the most influential brain regions (hubs) across all clusters contributing to each direction. Notably, the top-10 hubs with the largest proportional contributions were predominantly left-lateralized, specifically in clusters where the N group exhibited higher dwell-time.

This pronounced left-hemisphere lateralization of key dFC clusters aligns remarkably with a large body of literature on the asymmetrical nature of AD pathology. Although our findings do not imply a direct causal link between dwell-times and AD pathophysiology, they provide compelling correlational evidence. It is well-documented that brain atrophy in AD often develops unevenly, disrupting normal brain asymmetry (Friedland et al., 1988). Multiple studies have reported that cortical thinning, A*β* deposits, and neurite connectivity loss are more pronounced in the left hemisphere of AD patients (Lubben et al., 2021; Roe et al., 2021). This structural degradation is often accompanied by functional changes; for instance, a shift toward rightward asymmetry in white-matter networks has been interpreted as a consequence of left-hemisphere impairment in AD (Yang et al., 2017). Similarly, functional near-infrared spectroscopy (fNIRS) studies, which also measure hemodynamic responses, have observed reduced functional connectivity in AD patients, predominantly affecting left-hemisphere regions (Mizrak et al., 2024).

Our results contribute to this literature by demonstrating that dFC patterns may also reflect this left-hemisphere vulnerability in individuals at risk (in our study, this is an association with genetic factors) for AD. The identification of a distinct, significant, left-lateralized cluster (where dwell-time for N was higher) likely reflects the preservation of functional integrity in the not-at-risk group. This finding is consistent with the hypothesis of left-hemisphere vulnerability in AD (Bobkova and Vorobyov, 2015), though further research is needed to determine if this pattern represents a specific compensatory response.

Having established this group-level neurobiological association, we next asked whether these dFC-derived dwell-time features also contained sufficient information for individual-level genotype classification. Our initial machine learning models supported this possibility, yielding modest but consistently above-chance classification performance (Table 4). Importantly, the resulting models were interpretable, and when we summarized logistic regression coefficients by dwell-time direction, we observed a consistent pattern where increased dwell-time in clusters that were more characteristic of the N group was associated with reduced odds of risk-group membership (OR *<* 1), whereas increased dwell-time in clusters more characteristic of the risk groups was associated with increased odds of risk-group membership (OR *>* 1; Table 5). However, because overall predictive performance was moderate (69.6% AUC for N vs A+P- and A+P+), we evaluated whether the observed AUC values were statistically robust or could arise by chance using a non-parametric permutation framework (1000 permutations), where observed AUC was compared against a null distribution from models trained on randomly permuted genotype labels.

The permutation analysis provided evidence that dwell-time features carry a reliable genotype-related signal for specific contrasts (Table 6). In particular, classification was statistically significant for N vs. A+P+ (AUC = 0.680, *p* = 0.034) and for the pooled contrast N vs. (A+P-& A+P+) (AUC = 0.696, *p* = 0.005). Conversely, N vs. A+P-showed a trend-level effect that did not reach the conventional threshold (AUC = 0.636, *p* = 0.067), and A+P-vs. A+P+ was consistent with chance performance (AUC = 0.479, *p* = 0.486). Taken together, these results suggest that dwell-time differences are most reliably detected when contrasting the non-carrier group (N) against risk-group phenotypes, particularly the combined risk groups and the A+P+ subgroup, while differences between the two risk groups themselves are not well-captured by dwell-time features alone. This pattern is consistent with the interpretation that APOE-*ε*4 carriage (A+) may be the stronger driver of separability from the N group in our dataset than PICALM rs3851179 status (P+) alone, although larger cohorts will be required to confirm this more definitively.

Finally, although we believe that the current research makes novel contributions to the study of AD, our research does have limitations that must be addressed in future studies. First, the sample size is modest (68 subjects), which limits statistical power and increases uncertainty in both model estimates and permutation-based inference. Second, our current framework relies only on rs-fMRI. Structural MRI measures (e.g., regional volumes and cortical thickness) could capture complementary anatomical differences, and EEG could add sensitivity to fast neural dynamics. In addition, future work could extend this framework with mediation analyses to test whether neuroimaging-derived features statistically mediate relationships between genetic risk and downstream cognitive or behavioral measures. However, these additional analyses are beyond the scope of the current study. Third, our study remains an association study: we observe relationships between dFC-derived dwell-time patterns and genotype groups, but these findings should not be interpreted as causal. Despite these limitations, the convergent evidence from hub patterns and permutation-supported classification suggests that dFC dwell-time captures meaningful genotype-related variation. A key next step is validation in larger and ideally longitudinal cohorts, which will be necessary to evaluate stability over time and to better characterize preclinical AD risk signatures.

Overall, our results provide a direction for future preclinical AD detection studies by using dFC-derived features to reveal novel associative markers that may support earlier identification of AD risk.

## Conclusion

Our study used dFC with k-means clustering to extract dwell-time features that reflect how long individuals remain in recurring connectivity patterns. We identified two directionally opposite cluster types that differentiated the non-carrier group (N) from the risk groups (A+P- and A+P+): one in which the N group showed higher dwell-time than the risk groups, and another showing the opposite pattern. These effects were consistent across *k* = 4–9. Hubness analysis further showed a left-hemisphere bias among the top hubs in clusters where the N group had higher dwell-time, aligning with prior reports of asymmetric vulnerability in AD patients. At the individual level, logistic regression models achieved modest but above-chance classification performance. Importantly, permutation testing supported statistically significant discrimination for N vs. A+P+ and for N vs. the pooled risk groups, while N vs. A+P-showed a trend-level effect and A+P-vs. A+P+ remained at chance. Overall, these findings provide correlational evidence that dFC dwell-time patterns capture subtle genotype-related variation. These results motivate follow-up validation in larger and ideally longitudinal cohorts to assess stability over time and relevance to preclinical AD risk.

